# An experimental approach to study multi-species interactions implicated in Fusarium-Head Blight complex assembly in wheat

**DOI:** 10.1101/2024.04.14.589412

**Authors:** Miguel Ángel Corrales Gutiérrez, Harleen Kaur, Francesca Desiderio, Dimitar Kostadinov Douchkov, Jiasui Zhan, Heriberto Vélëz, Salim Bourras

## Abstract

Fusarium Head Blight (FHB) is a major disease of cereal crops present everywhere where wheat is produced. A complex of several, distinctly different, species from the genus *Fusarium* are responsible for causing the disease, with important taxa being *F. graminearum*, *F. culmorum*, and *F. avenaceum*. FHB is an adult stage, late season disease affecting wheat at anthesis. However, recent discoveries are now challenging the basic assumption that anthers and spikes are the only ecological niches relevant for resistance to FHB. Most prominently, it has been established that pathogens typically occurring earlier in the season, such as *Zymoseptoria tritici*, induce strong systemic changes in the plant immune system, that expand way beyond their primary ecological niche. Furthermore, taxa implicated in FHB can be isolated from virtually all wheat tissues, including typical blotch symptoms on wheat leaves. Strikingly, such important biological observations about the complex lifestyle of FHB causing taxa are overlooked as potential targets in resistance breeding. In this work we present a simplified approach allowing the dissection of the intricate lifestyles of the FHB causing *Fusarium* taxa. In doing so, we provide an experimental proof-of-concept that systemic signals from *Z. tritici* can affect the basic response of wheat to *F. graminearum*. We further demonstrate that differences in *Fusarium* species composition affect the outcome of the interaction in a genotype-specific manner. We argue that efforts to study hidden layers of the biology of FHB associated taxa are crucial to developing new resistance breeding strategies, based on a better understanding of the genetic drivers of the disease.

## Introduction

Fusarium Head Blight (FHB) is a wheat disease caused by a multispecies complex of up to 17 distinctly different *Fusarium* and non-*Fusarium* taxa (Karlsson *et al*., 2021). In nature, *Fusarium* ssp. are ubiquitous colonizers of monocot grasses that can be found in all tissues from roots to seeds. This is consistent with the piling evidence suggesting that pathogenic *Fusarium* taxa have evolved from endophytic *Fusarium* species, including those implicated in FHB (Hill *et al*., 2022; Ma *et al*., 2013). Despite this evolutionary trajectory, the *Fusarium* genomes have no clear signatures of lifestyle transition, even though such events are frequent in *Fusarium* fungi (Hill *et al*., 2022). This is strikingly contrasting with other ubiquitous colonizers such as Hypocreales fungi, where comparative genomic studies have revealed clear, lifestyle specific, genetic signature underlying the ability of these fungi to inhabit different hosts and environments (Wu & Cox, 2021). This suggests that lifestyle requirements in *Fusarium* fungi are very plastic, which is indeed the case with those associated with FHB. For instance, *F. graminearum* can survive as a saprophyte on wheat debris, while the most common pathogenic taxa implicated in FHB including *F. graminearum*, *F. culmorum*, and *F. avenaceum* can be found as latent endophyte inhabiting wheat stems, leaves, and roots (Kaur & Vilvert, *et al*., 2024; Lidia *et al*., 2021). Furthermore, it was also shown that FHB associated taxa can cause blotch-like symptoms on wheat leaves thus demonstrating that lifestyles shifts can occur in tissues where *Fusarium* spp. are present as silent endophytic colonizers (Kaur & Vilvert, *et al*., 2024).

Altogether, these data indicate that FHB causing taxa are present in the wheat host, very early in the season, throughout development, prior to anthesis. This is major because the etiology of FHB is often assimilated to the biology of *F. graminearum* with a two-step process is initiated at anthesis by for instance airborne spores landing in the anthers, which further spread through the rachis. Several studies have addressed the relative predominance of the different taxa implicated in FHB in relation to years and/or location. In Australia, several FHB epidemics in the 1980s, 1990s, and 2000s were incited by *F. pseudograminearum* (Burgess *et al*., 1987; Obanor *et al*., 2013; Obanor & Chakraborty, 2014). This is striking, because *F. pseudograminearum* is mostly associated with a disease called Fusarium Crown Rot, which affects a distinctly different wheat tissue at a much earlier developmental stage (Kazan & Gardiner, 2018). Thus, there can be large variations in complex composition, including several cases where *F. graminearum* was not the most prevalent species (Doohan *et al*., 2003; Lukanowski *et al*., 2008; Parry *et al*., 1995; Waalwijk *et al*., 2003; Xu *et al*., 2005). For instance, a field study evaluated the percentage of FHB infected wheat ears colonised by *F. graminearum*, *F. culmorum*, *F. avenaceum*, *F. poae*, and *M. nivale* in seven location throughout east and west Flanders in Belgium (Audenaert *et al*., 2009). The authors found large variation in complex composition across wheat genotypes and environments, with several cases where *F. poae* was the most predominant species causing FHB (Audenaert *et al*., 2009). In Italy, a large population study showed large shifts in FHB complex composition where *F. graminearum* was replaced by *F. poae* as the main taxon, and even supplanted by *M. Nivale* and *F. verticillioides* (Valverde-Bogantes *et al*., 2020). Finally in Kenya, a field survey where the incidence and severity of FHB were assessed at the hard dough stage showed that *F. avenaceum*, *F. poae* and *F. tricinctum* were the most abundant taxa. In one case. *F. graminearum* was the lowest in abundance of all isolated taxa (Maina *et al*., 2016).

This again demonstrates the ecological plasticity of FHB associated taxa, further highlighting the importance of lifestyle transition as a potential driver of FHB, and strengthening the argument that resistance to this disease cannot be reduced to *F. graminearum*. It also challenges to genetic assumptions restricting the definition of resistance to FHB to resistance to initial infection (Type I), resistance to the spread of the fungus through the rachis (Type II), resistance to toxin accumulation or the ability to degrade the mycotoxins (Type III), resistance to kernel infection (Type IV), and finally tolerance to yield loss (Type V). Strikingly, all types are restricted to flower/spike as the only tissue relevant to resistance, and all except ‘Type IV’ have a strong developmental stage-dependent, structural component. Furthermore, ‘Type IV’ is strongly challenged by recurrent and repeated observations that associations between mycotoxin production and FHB severity are inconsistent, which is a reasonable expectation considering piling evidence that *Fusarium* mycotoxins as antimicrobial compounds primarily directed against other microbes (i.e. not the host) (Sweany *et al*., 2022). Furthermore, *Fusarium* mycotoxin production in wheat is possibly a reminiscence of ancient association with grasses (i.e. the wild ancestors of cereals) where such compounds provide protection against herbivores (Gerling *et al*., 2023). Indeed, wild grasses serve as reservoir for *Fusarium* infections and mycotoxin contamination of wheat, which pinpoints the importance of understanding the endophytic lifestyle and its plasticity among members of the FHB species complex (Gerling *et al*., 2023).

Thus, the very genetic models serving as a basis for resistance breeding fail at accounting for such complexity, and the interplay between different members of the FHB complex. Similarly, such models also overlook important findings highlighting the importance of pathogen-induced systemic signals to wheat immunity (Seybold *et al*., 2020). It has been shown that infection with the foliar wheat pathogen *Zymospetoria tritici* induces systemic and long-lasting immunosuppression in the host (Seybold *et al*., 2020). Such effect was observed in distant organs and tissues that were not infected by *Z. tritici*, thus suggesting that early-stage infection of wheat with foliar pathogens can alter the response to disease occurring later in the season with the most prominent example being FHB. Another striking observation challenging the resistance ‘Type’ nomenclature, is that none of the 556 QTL identified in wheat as conferring resistance to FHB correspond to a major locus with dominant effect (Steiner *et al*., 2017; Venske *et al*., 2019). Furthermore, not a single immune receptor has been shown to control any form of resistance to FHB (Steiner *et al*., 2017; Venske *et al*., 2019). This is striking considering that the exceptional ability of *Fusarium* spp. to colonize a variety of tissues as silent apoplectic endophytes, which requires initial and permanent suppression of pattern recognition receptors (PRRs) (Saijo *et al*., 2018; Stotz *et al*., 2014). Indeed, the *Fusarium* genomes encode effector proteins, which is a class of fungal factor commonly associated with the suppression of plant immunity in fungal pathogens and non-pathogenic fungal endophytes (Collinge *et al*., 2022; Redkar *et al*., 2022; Rovenich *et al*., 2014). Furthermore, prominent members of the FHB complex such as *F. graminearum*, and *F. culmorum* behave as typical hemibiotrophs in leaf infection assays, with an initial latent, asymptomatic biotrophic phase followed by necrosis induction (Diamond & Cooke, 1999; Kaur & Vilvert, *et al*., 2024).

Thus, the basic expectation is that the wheat immune response to FHB should involve some form of receptor-ligand interaction implicating well defined host immune receptors. However, the ability to genetically explore such hypothesis is largely restricted by the phenotyping strategies used to score FHB, which are largely aligned to the pre-defined resistance types I to V. Most prominently, interactions between FHB causing taxa and the host are almost exclusively assessed from anthesis onwards, with a large focus on scoring blight symptoms on the spikes and mycotoxin accumulation in the grain (Alisaac & Mahlein, 2023; Khan *et al*., 2020; Mesterhazy, 2020). Still, several alternative phenotyping strategies have been developed to allow large phenotypic screens of more complex traits that cannot be implemented in the field. These include detached leaf assays (Diamond & Cooke, 1999), seedling resistance assays (Mesterhazy, 1987), seed germination assay (Browne & Cooke, 2005), and coleoptile infection assay (Wu *et al*., 2005). Strikingly, while none of these strategies is targeting anthesis, all have proven useful for the identification of resistance traits against FHB in wheat and other cereals. Most prominently, and of direct relevance to this work, assays based on artificial infections of detached leaf assay has been particularly used due to the high correlation between pathogen latent period and quantitative disease resistance in the field (Diamond & Cooke, 1999; Niks & Skinnes, 1998).

In this work we present a phenotyping strategy designed to investigate the host genetic factors affecting the endophytic lifestyle of *Fusarium* taxa associated with FHB and their ability to associate in a complex. We developed a fully featured phenotyping strategy based on highly controlled detached leaves infections, combined with machine-learning aided image analysis allowing accurate quantification of disease symptoms. In doing so, we also produce an experimental proof-of-concept for the potential role of systemic signals and the host genetic background in controlling lifestyle transition and complex assembly, thus providing a working system for mining potentially novel FHB resistance traits.

## Methods

### Plant growth conditions

An overview of the phenotyping pipeline is provided in **Figure 1**. In this study, we used the reference hexaploid wheat cultivars Chinese Spring, Fielder, and Bob-white; the breeding lines Agadir, Artico, and Victo which are *Triticum aestivum* spp. *aestivum*; Latino, a *T. turgidum* spp. *durum* cultivar, MG5323, a *T. dicoccum* accession, and Zardak, an old cultivar of *T. turgidum* spp. *turanicum*. We also used 30 commercial spring wheat cultivars commonly grown in Sweden provided by the Swedish Agricultural Cooperative Lantmännen Lantbruk (Sweden). All genotypes were germinated in pots with sterilized substrate at 20-21 °C, RH of 70% and with light intensity of 300 μmol and 18 h light/ 6 h dark photoperiod. Plants for inoculation with *Z. tritici* were maintained in a closed infection chamber inside a growth chamber with the same above conditions, and supplemented with additional LED light sources enriching the light spectrum in the 380 and 700 nM wavelength, respectively (**Supplementary Figure 1**).

**Figure 1.**
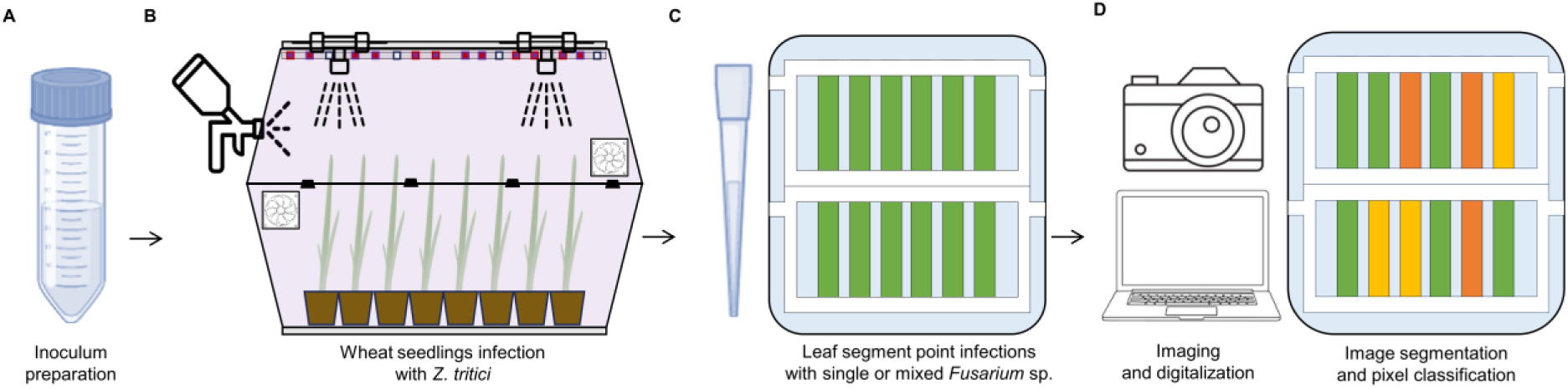
Overview of the phenotyping pipeline. A 11-days-old *Z. tritici* (an hemibiotroph) culture was scratched from YMS plates, resuspended in TWEEN 0.01% (A) and sprayed in a closed environment (named as The Hive) where plants have kept until the appropriate stage (B). After 3 days, leaf fragments are harvested, mounted on agar plates and infected with a spore suspension of *Fusarium* spp. (a necrotroph) (C). After 4-7 dpi of incubation, pictures of the disease symptoms were taken with a Nikon camera. The pictures were analyzed with bash code throughout python scrIpts and the results were analyzed with R (D).

### Preparation of *Zymoseptoria tritici* inoculum

*Z. tritici* isolates ST99CH_3B8 (abbreviated as 3B8), ST99CH_3C4 (3C4), ST99CH_3C7 (3C7), ST99CH_3D5 (3D5), ST99CH_3D7 (3D7), ST99CH_3F2 (3F2), ST99CH_3G6 (3G6) were isolated in Switzerland in 1999 (courtesy of Prof. Bruce McDonald, Federal Institute of Technology, ETH Zürich, Switzerland). *Z. tritici* isolates were incubated in yeast extract-malt extract-starch agar medium (YMS 4 g/L yeast extract, 4 g/L malt extract, 4 g/L sucrose, 16 g/L agar), in darkness at 20 °C for 11 days until the day of the infection. Spores were collected from the plates by scraping the cells in sterile conditions and resuspend in TWEEN 0.01%. The solution was filtered with a cloth with pore diameter of 22-25 µm. Spore concentration was determined with a hemocytometer and adjusted in all isolates to 1×10^6^ spores/ml. Before air gun infection, all isolates were mixed at equal volume and concentration.

### Preparation of *Fusarium* inoculum

*F. avenaceum*, *F. culmorum* and *F. graminearum* were collected from Swedish fields in summer of 2021 and characterized morphologically for species determination, and genotyped by Kaur & Vilvert *et al*. (2024) using universal ITS primers ITS1 (5’-TCCGTAGGTGAACCTGCGG-3’) and ITS4 (5’-TCCTCCGCTTATTGATATGC-3’), and species specific primers for *F. graminearum* marker GOFW (5’-ACCTCTGTTGTTCTTCCAGACGG-3’) GORV (5’-CTGGTCAGTATTAACCGTGTGTG-3’), and *F. culmorum* with marker Fc03 (5’-TTCTTGCTAGGGTTGAGGATG-3’) Fc02 (5’-GACCTTGACTTTGAGCTTCTTG-3’) (Astrid Bauer & Seigner, 2015; De Biazio *et al*., 2008). For co-infections with the three *Fusarium* species, isolates were grown for two weeks at 25 °C on oatmeal agar (OMA: 3 g/L organic oatmeal flower and 16 g/L agar) and sporulation was induced by exposure to natural light for 3 days. For co-infections with *Z. tritici*, all *Fusarium* species were incubated in muesli agar (MA: 25 g/L organic muesli: oatmeal, wheat spikes, barley and rye flakes, sunflower seeds, and sunflower lecithin; 2 g/L malt extract and 12 g/L agar), and incubated at 25 °C for 30 days. After the incubation period, spores were collected by scratching from the plate and suspended in TWEEN 0.01%. The final spore concentration was adjusted to 250 spores/μL and aliquoted into vials at −20 °C until infection day.

### Coinfection assays

For coinfections with different *Fusarium* species, 6 cm long leaf segments were cut from 15-days-old wheat seedlings. Leaf segments were punctured and inoculated with 15 μL of a spore solution at the desired concentration. For co-infections with different *Fusarium* species, an equal volume and concentration of spores (10 spores/μL) were mixed into a 15 μL drop and inoculated as described above. As a control, a group of leaf segments were inoculated with TWEEN 0.01%. Leaf segments were placed in Benzimidazole Agar Medium (BAM: 5% water agar supplemented with 0.03 mg/L benzimidazole). Custom frames made of polylactic acid (PLA) were designed and 3D printed on a Flashforge Guider 2s machine (Flashforge). These were used to keep the leaf surface in contact with BAM medium, preventing desiccation and curling which happens when incubation time extends over 4-5 days. Pictures were taken at 8 dpi using a DSLR camera. Phenotyping of the commercial cultivars, lab standards, and breeding lines was performed using shorter 4 cm length leaf segments, with a spore concentration of 250 spores/μL and an incubation time of 5 days without PLA frames.

For *F. graminearum* – *Z. tritici* coinfections, 9-days-old wheat seedlings were placed in closed chamber made of two stacked clear PLA containers. The *Z. tritici* inoculum consisted of a mix at an equal ratio of the seven isolates described above, sprayed with a spray gun over the seedlings using 0.33 ml per seedling. As a control, a group of plants were inoculated with TWEEN 0.01%. After inoculation, the climate in the infection chamber was maintained at high humidity (over 70% of RH) and in darkness for 24 hours. Then plants were placed back in a day-night cycle as described above, and kept at high RH by spraying distilled water, using a low-pressure spray gun. Then, 6 cm length leaf segments were cut from *Z. tritici*-infected and mock-inoculated plants at 3 days post infection (dpi). Secondary infections with *F. graminearum* were performed as described above, using a 15 μL drop with a *F. graminearum* spore concentration of 250 sp/μl. Leaf segments were kept on BAM plates and maintained in darkness with high RH for 24 hours. Then, plates were moved back to a day/night cycle at high RH, incubated for 4 days (MG5323) or 5 days (Fielder), then photographed.

### Machine learning-aided image analysis

Each leaf segment from the original picture was extracted individually using ImageJ. To quantify disease severity, Jupyter Notebooks were used to run code from the PlantCV and OpenCV packages (Gehan *et al*., 2017). Images showing the juxtaposition of different masks were obtained using the Jupyter Notebooks terminal interface. Bash commands were used to simultaneously analyze multiple samples from one directory with the PlantCV based script, which creates a JSON file as output for further analysis. The implementation of PlantCV allows binary image transformation from original RGB pictures, mask creation, and pixel classification in different predefined classes using a multiclass Naïve Bayes (NB) algorithm.

To define our region of interest (ROI) and reduce the error rate of the algorithm, the picture area inside the PLA frames holding the leaves was extracted. Each leaf segment was extracted as individual image using PlantCV based script (Gehan *et al*., 2017). Pixels were manually captured with ImageJ, and their RGB parameters (red, green, blue) were saved in a tab delimited file where we define each class of pixel. To avoid class imbalance, we collected the same number of pixels for each class (necrosis, chlorosis, healthy tissue, and background). The training set consisted of over 1000 different pixels from each class. Once the RGB values of each class were defined in a training set, a Naïve Bayes (NB) algorithm (Gehan *et al*., 2017) was used to develop a probability function for each pre-defined class. After the training step, a probability function was used by the NB algorithm to classify each pixel, and the data was exported to JSON file containing the number of counted pixels in each predefined class. Data was transformed to a tabular file and uploaded to the R statistical environment for further analysis.

### Statistical analysis

Data was initially tested for normality and homocedasticity, then a linear model with fixed effects was designed after performing an aligned-rank transformation with the values of necrosis or chlorosis (*y*) as response variable and cultivar (*x*_1_), zymoseptoria (*x*_2_) and its interaction (*x*_3_) as explanatory variables with their corresponding coefficients (*β*_1_, *β*_2_, *β*_3_). The constant µ represents the intercept.

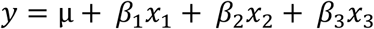

To test statistical differences, we performed an aligned-rank-transformation-two-way-ANOVA (robust two-way-ANOVA) and a *post hoc* test with Tukey adjustment. A Principal Component Analysis (PCA) was carried out with the average value of each cultivar and *Fusarium* species combination (each combination has at least 5 biological replicates). Coordinates of each sample in the PCA were extracted to calculate the distance matrix which was used as input for cluster analysis using a Hierarchical Agglomerative Nesting (AGNES) algorithm. All statistical analyses were performed with R (version 4.2.2) in the Rstudio environment (R Core Team, 2022) and the packages: readxl, dplyr, tidyr, stringr, reshape2, ggplot2, multcompView, forcats, ARTool, rcompanion, factoextra, and cluster (Graves *et al*., 2023; Kassambara & Mundt, 2020; Kay *et al*., 2021; Maechler *et al*., 2022; Mangiafico, 2023; Wickham, *et al*., 2023; Wickham, *et al*., 2023; Wickham, 2007, 2016, 2022, 2023; Wickham & Bryan, 2023).

## Results

### Phenotypic analysis of single *vs.* mixed *Fusarium* species infection of wheat leaves

We selected the bread wheat cultivar Fielder for this assay based on the consistent and clear phenotypes we obtained in our preliminary tests and our previous work (Kaur and Vilvert, *et al*. 2024). We adapted a leaf segment infection strategy based on previous work demonstrating high correlation between the phenotypes observed in leaf segment assays and quantitative disease resistance to FHB in the field (Kumar et al., 2011; Perochon & Doohan, 2016). Leaf segments were inoculated with *F. avenaceum*, *F. culmorum*, *F. graminearum* and their combinations by drop inoculation (see Methods). At 8 dpi, variability in disease severity among the different pathogens and their combinations was assessed by eye (**Figure 2**). Here, we reasoned that if interactions between *Fusarium* species are a major component of how these species act in a complex, then such assay should lead to clear differential phenotypes i.e. observable by a trained person. Indeed, *F. graminearum* showed the most severe symptoms, triggering necrosis in large areas of the leaf segment. By contrast, the combination of *F. culmorum* and *F. graminearum* caused much weaker symptoms as compared to single infections with *F. graminearum*. These results suggest that *F. graminearum* is the most aggressive of the three *Fusarium* species tested, and that there are interactions among *F. avenaceum*, *F. culmorum* and *F. graminearum* that could change the plant response and or directly affect complex assembly. We conclude that the detached leaf assay is sensitive to *Fusarium* species composition and can reveal quantitative difference in disease severity resulting from pure *vs*. mixed species inocula.

**Figure 2.**
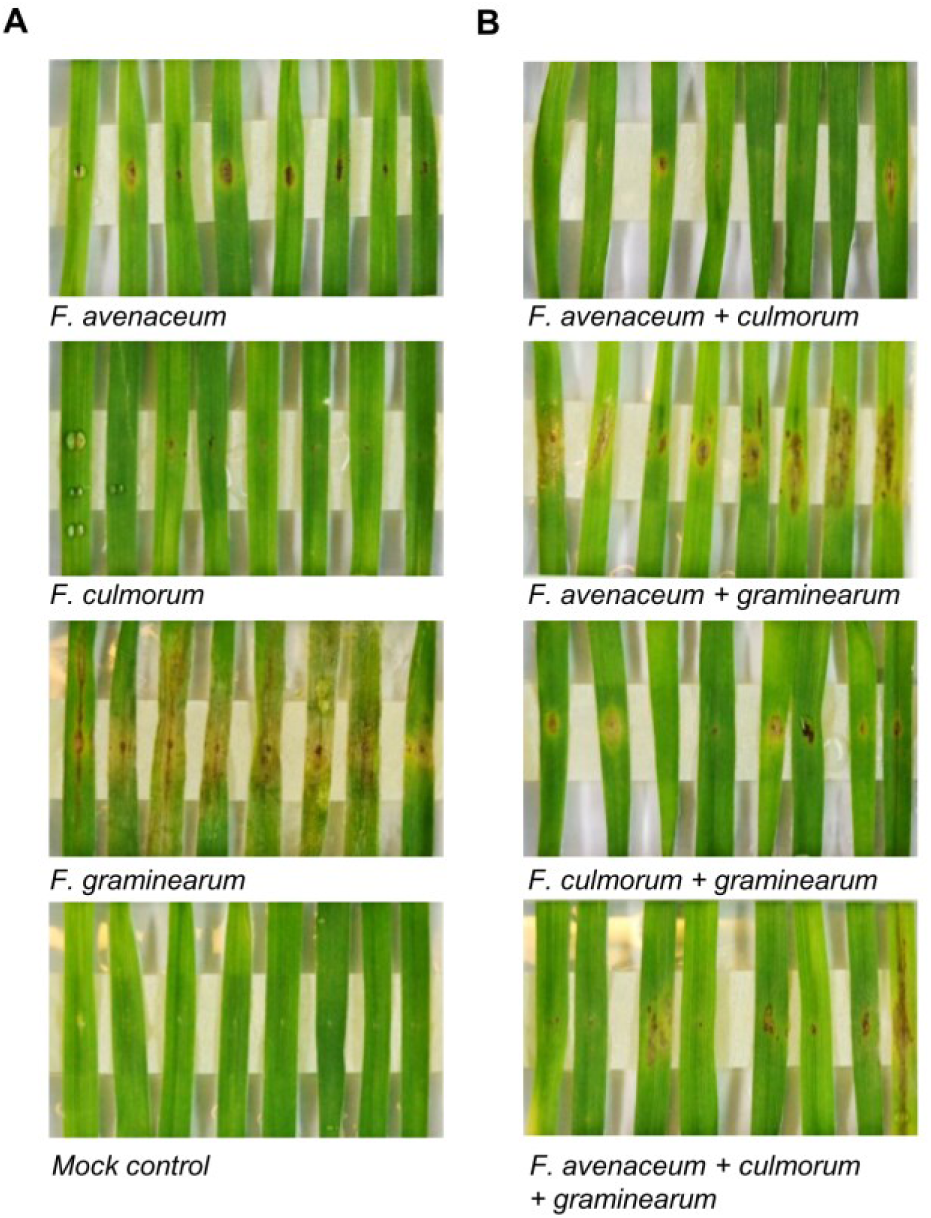
Phenotyping of Fusarium coinfections. The cultivar Fielder was inoculated with *F. avenaceum*, *F. culmorum* and *F. graminearum* isolates from Sweden. Pictures were taken at 8 dpi. We perform two independent experiments with 8 biological replicates per treatment. We show representative pictures of one of the experiments.

### Precision phenotyping using Image-based quantification of disease severity

In another series of experiments, we aimed at developing an image-based precision phenotyping strategy to distinguish and quantify two important components of the observed symptoms, namely: necrosis and chlorosis. We reasoned that these two types of tissue alterations denote different physiological processes taking place during the progression of disease, and their prevalence may vary between genotypes thus pinpointing to distinctly different host factors involved in the response to *Fusarium* sp. We analyzed the pictures using a NB algorithm trained with a set of RGB values from different pixels classified as ‘necrosis’, ‘chlorosis’, ‘healthy tissue’ and ‘background’ (see Methods). We used a test set of 64 leaf segment pictures to test this approach. We were able to clearly separate each type of tissue, and create so called ‘masks’ (see Methods) allowing the visualization and the quantification of the infected area undergoing necrosis *vs*. chlorosis (**Figure 3**). The results of the quantification of each leaf segment grouped by treatment are illustrated in **Figure 4**. They also could validate that the script correctly identified the infection with *F. graminearum* as the infection which triggers the most severe response (the highest amount of necrotic tissue). The levels of necrosis observed in the control group are close to 0, indicating a high performance of the classifier algorithm. We could also corroborate and quantify the cases where mixed infection resulted in less necrosis than infection with a pure culture of *F. graminearum*. Altogether, we conclude that our image-based, machine learning aided phenotyping pipelines can correctly and precisely quantify disease symptoms and reveal the smallest differences in the wheat response to single *vs*. mixed *Fusarium* inocula.

**Figure 3.**
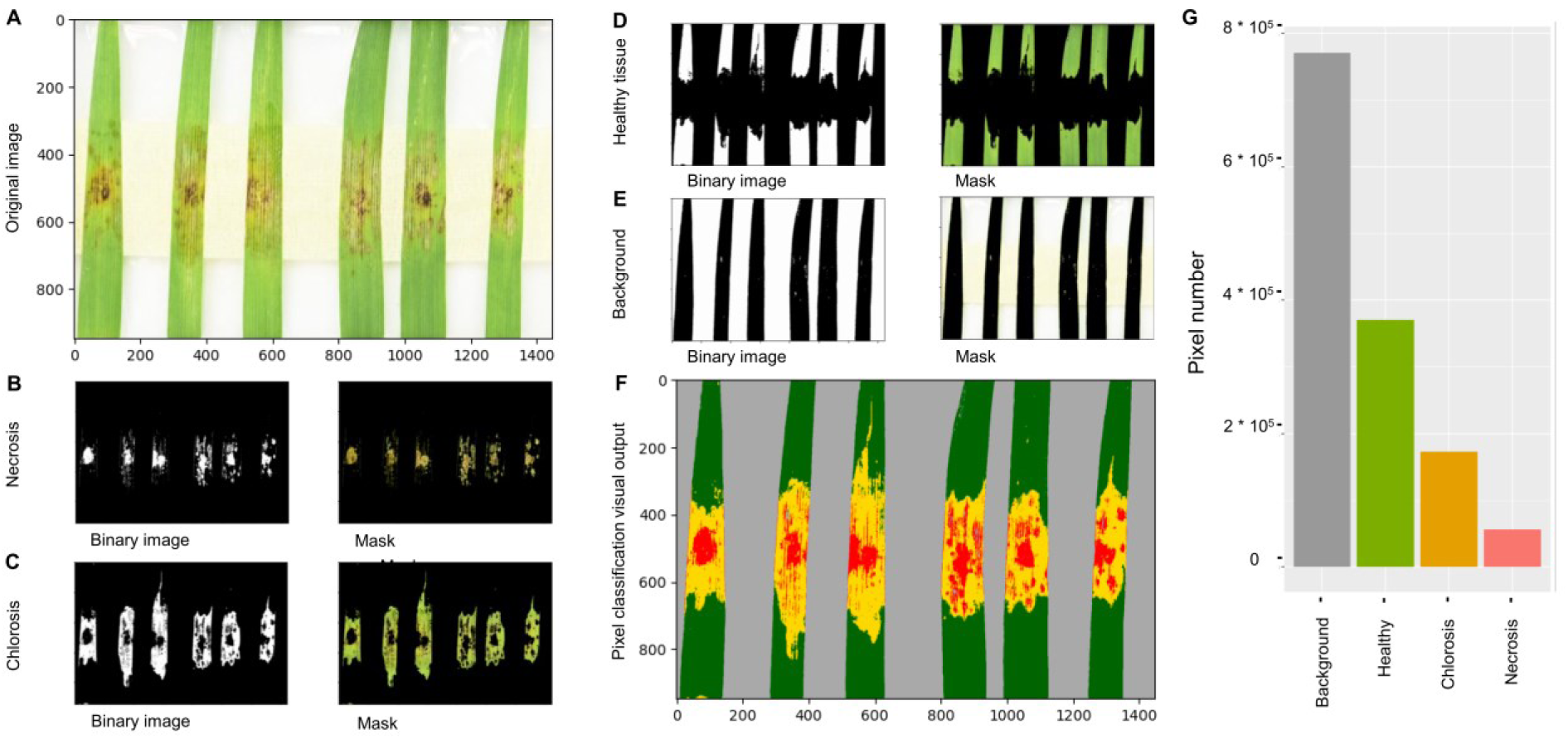
Steps for image analysis throughout quantification of Fusarium symptoms. A Naïve Bayes algorithm use a training set of pixels collected with the free software ImageJ to develops a probability function with the aim of classify all picture pixels in each of the predefined classes (necrosis, chlorosis, healthy tissue, or background). The result can be visualized in a binary image that is used to create a mask. All masks can be overlapped in a final image where is counted the number of pixels in each class, allowing the quantification of the disease symptoms.

**Figure 4.**
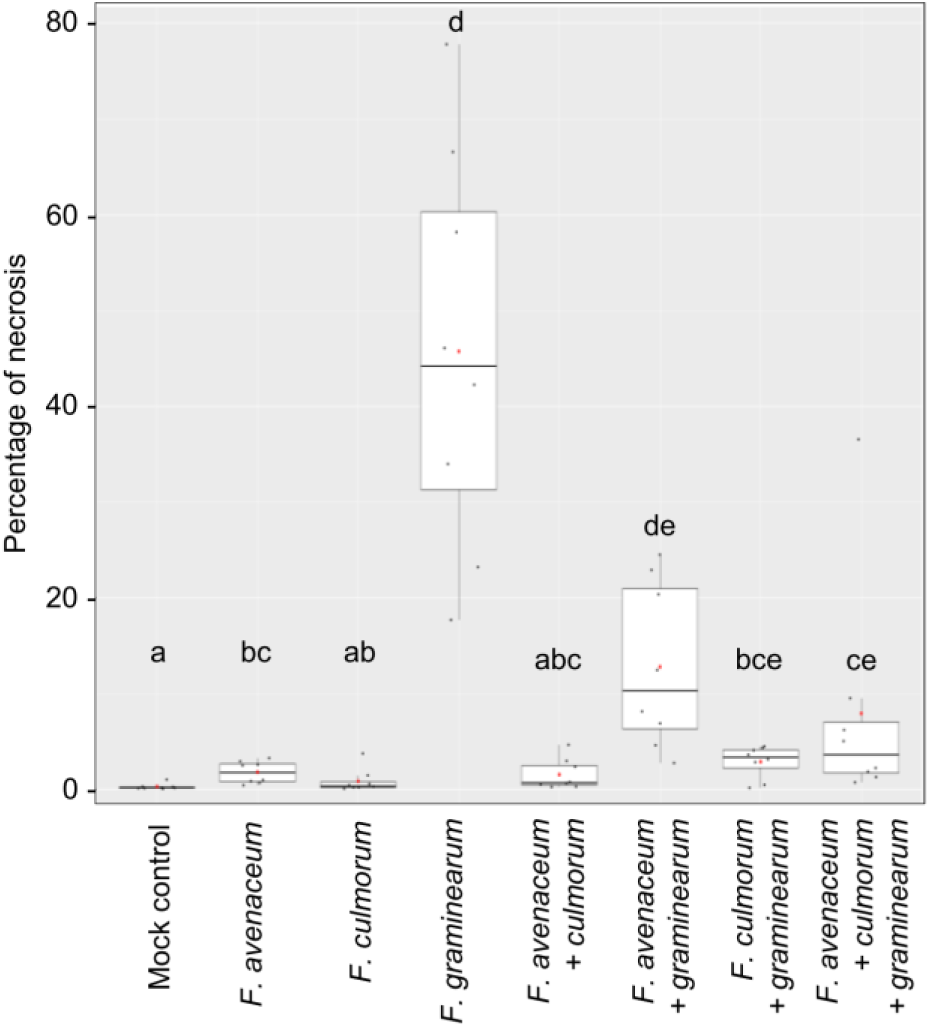
Symptom disease severity quantification through Naïve Bayes algorithm. Pictures from Figure 2 were analyzed by the set up shown in Figure 3. The number of pixels classified as “Necrosis” was divided by the number of pixels calculated in the different groups of leaf pixels (“necrosis”, “chlorosis” and “healthy tissue”). The letters a-e refer to two groups of treatments that present differences statistically significative (two-way robust ANOVA: p < 0.05)

### Comparison of disease severity values scored visually *vs.* using machine-learning

We then compared the disease scoring obtained from a trained person with the results obtained with machine-learning (ML) aided image analysis. Here, we aimed at identifying the added value of accurate segmentation of the phenotype into necrosis and chlorosis (i.e. expressed as a percentage), which we argued are very difficult to separate from the composite disease phenotype scored by eye (expressed as a scale from 1 to 5). We used an image library previously generated by us (Kaur and Vilvert., *et al*, 2024). It consisted of images of detached leaf infection assays with *F. avenaceum*, *F. culmorum*, *F. graminearum* and their combinations in a panel of different wheat lines. Due to the large number of putative comparisons (39 lines and 7 different treatments) and considering the complexity of the data, we decided to perform an exploratory cluster analysis with the coordinates of each sample obtained in a PCA and compare the relative position of each cultivar.

We found that the most coherent clustering was obtained with the percentage of necrosis scored by ML based on a Naïve Bayes algorithm (**Figure 5, Supplementary Tables 2, 3, 4, 5**). This approximation aggregates the samples in the breeding lines (blue), lab standards (pink) and commercial lines (green, red and yellow). Moreover, the clusters obtained with the values of necrosis percentage are similar to the groups obtained with visual scores. In both cases there are two clear groups of commercial lines close to the lab standard and the breeding lines (blue and yellow in manual scoring and orange and yellow in ML scoring), and a third commercial line group which is less similar to the lab standards (green, ML scoring), which is split in two closely related groups in the manual scoring (green and pink). In contrast, with manual scoring, the parental lines are spread between the cluster of lab standards (Latino, Zardak and MG5323, tetraploid durum wheat) and one of the clusters of commercial lines (Artico, Victo and Agadir, hexaploid wheat). However, beyond these differences, the relative position of each line in comparison with the others remains relatively similar. For example, the lab standard Fielder is close to Chinese Spring, Bob-White, Eleven, Amulett, and Rogue in both cases, with manual scoring and with necrosis calculated with Naïve Bayes algorithm. In summary, we found that the ML-based scoring allowed better defined clusters differentiating the different genotypes used in this study. These results further indicate that the ML strategy surpassed the visual scoring in terms of (i) tissue segmentation accuracy, (ii) sensitivity to small differences, and (iii) precise quantification of different tissue types, all of which are of paramount importance for clustering genotypes based on similar disease responses.

**Figure 5.**
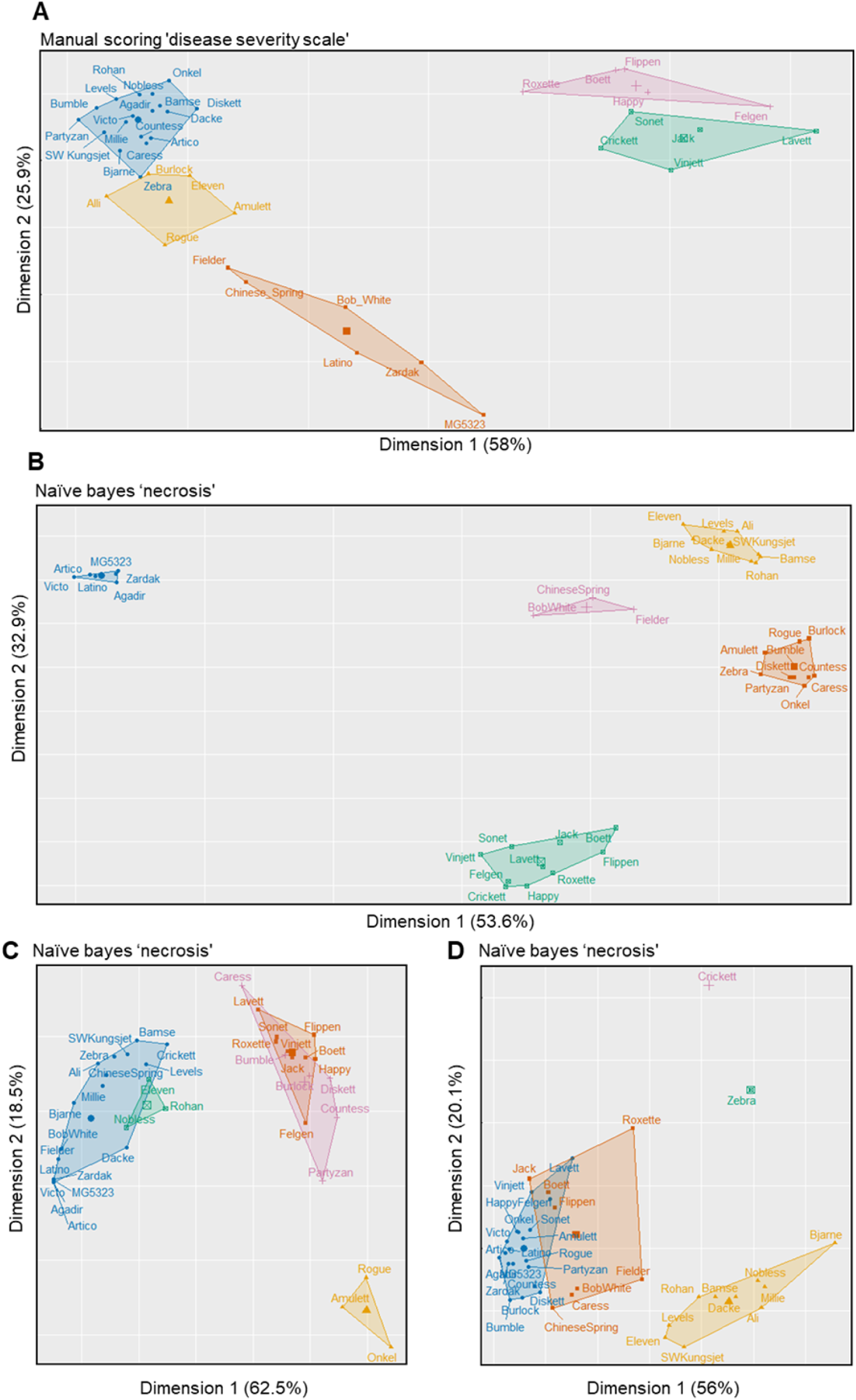
Cluster analysis of wheat cultivars infected with *F. avenaceum*, *F. culmorum*, *F. graminearum* alone or in combination. The Agglomerative nesting (AGNES) algorithm was applied to the distance matrix of the mean value of the PCA coordinates of each sample to develop the dendrogram. The defined number of clusters is 5 (k = 5).

We then performed a robust ANOVA analysis (**Supplementary Table 6**), to detect statistically significant differences between different groups of cultivars vs. treatments. The results of the ANOVA indicate that observed differences (i.e. as in the clustering) between the groups of cultivars, *Fusarium* combinations and the combination of both, are statistically significant (p < 0.01). This further demonstrates that our strategy can detect single *Fusarium* species and *Fusarium* complex-specific responses in a cultivar-dependent manner, which is an important validation of our strategy as a potential tool for mapping host factors underlying FHB complex assembly in wheat.

### Phenotypic analysis of the effect of primary infection with *Z. tritici* on *F. graminearum*

In a recent study by us we could demonstrate that *Fusarium* taxa commonly implicated in FHB largely co-exist with wheat leaves showing blotch symptoms typically caused by *Z. tritici* in Sweden (Kaur and Vilvert, *et al*, 2024). We further isolated several isolates of *F. graminearum* and *F. culmorum* from typical blotch-like necrotic lesions, and demonstrated using our leaf segment assays that they can indeed cause blotch-like symptoms that can be easily confound with *Z. tritici*, or *Pyrenophora-tritici repentis* (Kaur and Vilvert, *et al*, 2024). We thus aimed at developing a phenotyping strategy that would allow us to explore these intriguing interactions between *Z. tritici* and *Fusarium* spp.

We used cultivar Fielder, a universal susceptible control for *Z. tritici* which has also been the basis for our leaf segment infection assays with *F. graminearum*. We used a mix at equal ratio of seven *Z. tritici* isolates representing genotypes with a broad and non-overlapping spectrum of virulence originating from an extensively characterized population developed at ETH Zurich (Zhan *et al*., 2003, 2005). Considering the high genetic diversity of *Z. tritici* populations commonly present in a single field, we reasoned that our assays would gain in biological relevance may we integrate a higher-level of inoculum complexity. We also used *F. graminearum*, as it was the *Fusarium* specie which resulted in the most severe symptoms in our assay.

First, we checked the sensitivity of the image analysis strategy to detect dose-dependent effects where *F. graminearum* inoculations were done at 10, 100 and 250 spores/μl on material that was initially infected with *Z. tritici* (see Methods, **Figure 6A**). The NB algorithm recognized higher levels of necrosis and chlorosis in leaf segments infected at a concentration of 10 spores/μl in comparison with the negative controls, although the differences are not consistent enough to be statistically significant (**Figure 6B**). Leaf segments inoculated with a concentration of 100 and 250 spores/μl showed similar levels of necrosis and chlorosis, although higher concentrations tend to give more severe disease symptoms (**Figure 6B, 6C**). Most importantly, we also detected a reduction of the necrosis and chlorosis caused by *F. graminearum* on the leaf segments pre-infected with *Z. tritici* (**Figure 6B**). It is noteworthy that, considering the timing of the co-infections, the observed interactions are taking place during the biotrophic phase of the hemibiotrophic lifestyle of *Z. tritici*. Therefore, these results suggest there is possible antagonism from *Z. tritici*, during its biotrophic stage, to suppress the necrotic lifestyle of *F. graminearum*, which can be detected by our strategy.

**Figure 6.**
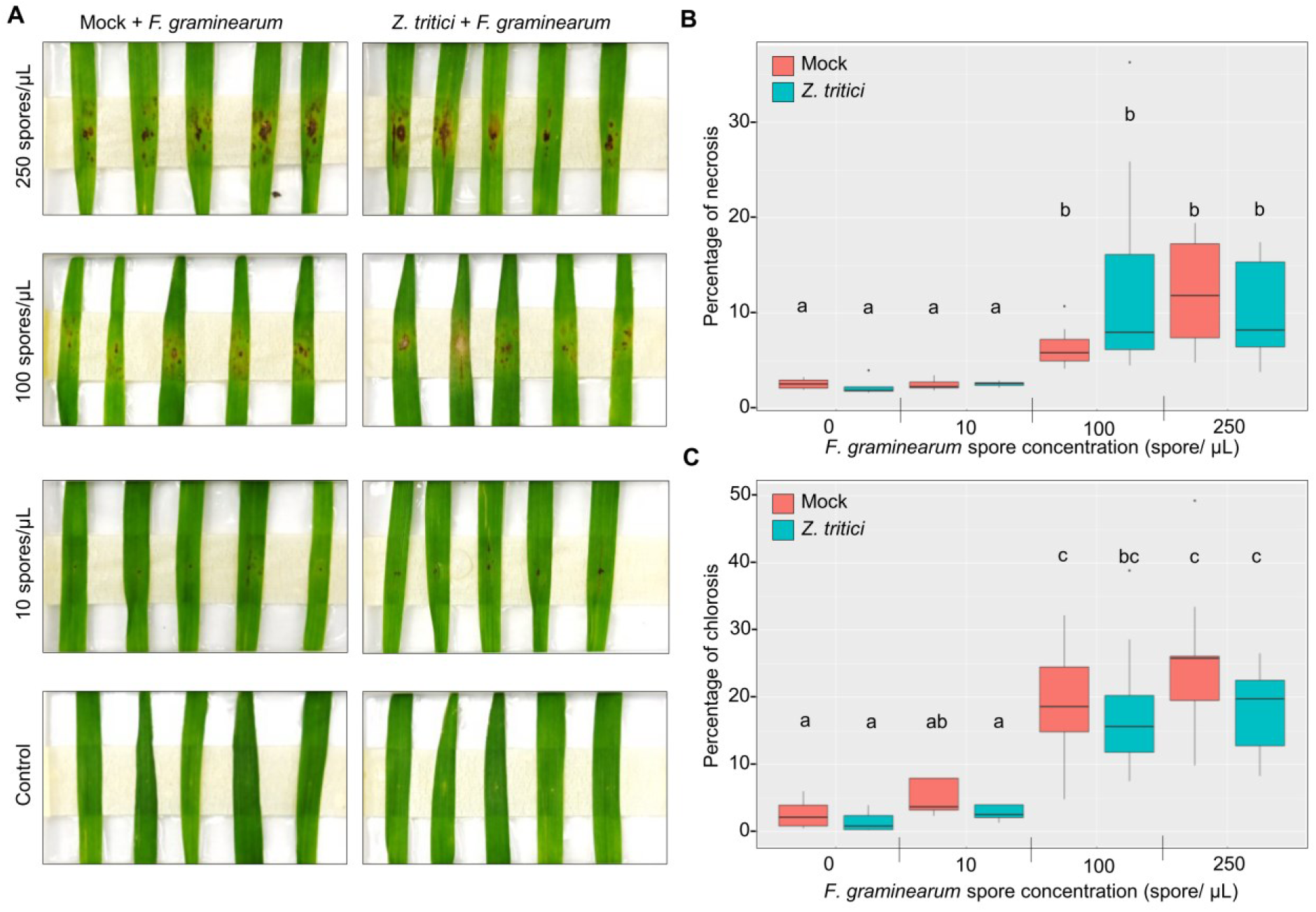
Quantification of the disease symptoms of *F. graminearum* at different spore concentrations in coinfections assays with *Z. tritici*. (A) Pictures from one experiment with 5 biological replicates were analyzed with our setup at 5 dpi. The symptom quantification was obtained dividing the number of pixels counted in the class necrosis (B) or chlorosis (C) between the sum of the number of pixels in necrosis, chlorosis and healthy tissue classes. The pink color indicates mock treatment and blue color indicates plants coinfected with Z. tritici. The letters a-b refer to two groups of treatments that present differences statistically significative (two-way robust ANOVA: p < 0.05)

To further explore the capacity of our pipeline to reveal such phenotypes, we expanded our analysis to one tetraploid wheat accession (MG5323) which has been consistently more susceptible to *F. graminearum* leaf infections than Fielder in our assays. We wanted to assess if an inhibitory effect from *Z. tritici* would still be visible provided with higher virulence from *F. graminearum* on the host. Here, comparisons between MG5323 and Fielder come handy as both genotypes are susceptible to *Z. tritici*, thus the outcome of interactions should be mainly influenced by the virulence level of *F. graminearum*.

The results from the image analysis show that our pipeline can still detect an inhibitory effect of *Z. tritici* on *F. graminaerum* (**Figure 7A**), however, this effect is weaker in MG5323 where *F. graminearum* is more virulent (**Figure 7B**). We also observed that such an effect seems to preferentially act on necrosis rather than chlorosis in Fielder, where inhibition allows more refined phenotypic readout. On the contrary, the effect on chlorosis was more prominent in MG5323 where we were able to detect a statistically significant reduction in presence of *Z. tritici*. Together, these results suggest that *Z. tritici* can modify the host response to *F. graminearum* which seems to be conserved across different wheat species. We propose that such mechanisms are likely to alter the endophytic lifestyle of the *Fusarium* species thus playing a possible role in shaping *Fusarium* populations in the field.

**Figure 7.**
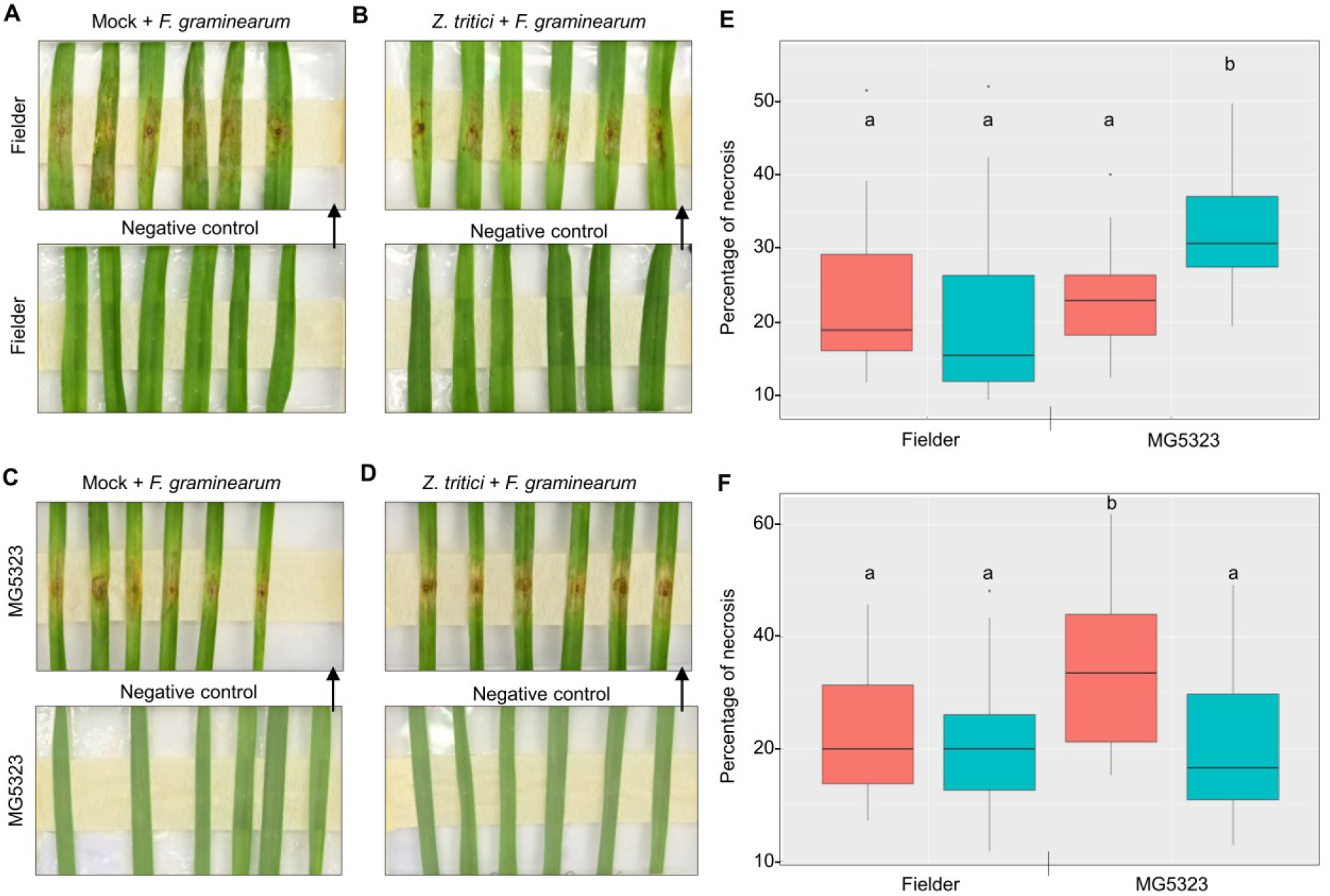
Quantification of the disease symptoms of *F. graminearum* in coinfections assays with *Z. tritici*. (A) Pictures from two experiments with 6 biological replicates were analyzed with our setup. We show characteristic pictures of one experiment. The symptom quantification was obtained dividing the number of pixels counted in the class necrosis (B) or chlorosis (C) between the sum of the number of pixels in necrosis, chlorosis, and healthy tissue classes. The pink color indicates mock treatment and blue color indicates plants coinfected with *Z. tritici*. The letters a-b refer to two groups of treatments that present differences statistically significative (two-way robust ANOVA: p < 0.05)

## Discussion

In this study we presented an original experimental set up allowing the study of disease interactions in wheat with a focus on FHB. Detached leaf assays are highly suitable for large scale screens due to the relatively short time from seed to inoculation, and the low requirements of space. In fact, the full procedure can be performed in 17 days thus facilitating the screening of large populations. Several procedures have been developed to phenotype resistance to FHB ranging from spikelet infection in the lab and in the field (Chhabra *et al*., 2024; Giancaspro *et al*., 2016), *in vitro* detached head inoculation (Huang *et al*., 2020), seedling assays (Soresi *et al*., 2015), coleoptile infection assays (Wu *et al*., 2005), and detached leaf assay (Perochon & Doohan, 2016). Even though detached leaf assays are commonly used, precision phenomics strategies to quantify the resulting phenotypes have not been exploited, with a few attempts restricted to the measurement of lesion diameter (Perochon & Doohan, 2016; Roesler *et al*., 2022). A major feature of our strategy is the possibility to simultaneously decompose the symptoms into two types of alterations (necrosis and chlorosis) and precisely (i.e. to the pixel) measure their relative areas. Furthermore, we demonstrate that there are measurable differences in the ability of FHB associated taxa to cause either of these symptoms on different genotypes (alone or in combination). Thus, in this work, we took full advantage of such reductionist approaches to explore the complexity of multi-pathogen species interactions within the FHB complex, and between FHB and other wheat pathogens.

The first interesting finding of this work was the inherent variation in the capacity of different FHB associated *Fusarium* species to induce chlorosis or necrosis. While the mechanisms underlying the expression of these tissue alterations are conserved among plants, their physiological basis is very different. Chlorosis is typically a nutrient deficiency phenotype commonly induced when micronutrients such as iron (Fe) and zinc (Zn) are depleted from the cells (Nogiya *et al*., 2016). Chlorosis is not to be confused with ‘senescence’ which is a highly regulated developmental process in plants which may also result from nutrient deficiency (i.e. starvation) but is largely controlled by genetic programming (Noodén *et al*., 1997). Necrosis is a typical marker for biotic stress, triggered by the plant immune system in response to pathogens (Minina *et al*., 2005). In this context, our results indicated that FHB associated species have different abilities to differentially alter two highly conserved tissue damage-associated mechanisms reminiscent of blight symptoms. This could indicate a form of functional specialization among species of the FHB complex.

Furthermore, we showed that the ability to cause these symptoms and their relative prevalence is genotype-specific, and species combination-dependent. Such genotype-specific variation in the hemibiotrophic lifestyle was also observed in other pathogenic *Fusarium* species such as *F. temperatum*, and *F. oxysporum* (Hill *et al*., 2022; Robles-Barrios *et al*., 2022; Srivastava *et al*., 2024). We therefore speculate that such variation may reflect dynamic changes in the hemibiotrophic lifestyle of *Fusarium* sp. Here, a classical progression would correspond to an early endophytic colonization without visible symptoms, followed by the induction of chlorosis as nutrients are gradually depleted by the fungus, then a shift to a necrotrophic lifestyle where the pathogen switches to feeding on dead tissues. Thus, what our assay could be revealing a possibly new types of resistance to FHB based on the suppression of deleterious tissue alterations by chlorosis and necrosis.

Another interesting finding from this study was the observation that co-existence of different *Fusarium* species can have a measurable negative impact on the performance of the complex to cause lesions on the leaves. In fact, we found a statistically significant difference between single *F. graminearum* infections and inoculations with *F. culmorum* and *F. graminearum*. This is counter intuitive considering that disease complexes such as FHB are largely considered as based on a form of synergism between microbial pathogens (Lamichhane & Venturi, 2015). This is for instance represented in our assays by the *F. avenaceum* and *F. graminearum* combination, which tends to have a higher disease severity than *F. avenaceum* single infection. Similarly, evidence shows that co-infection with *F. aglaonematis sp. nov.* and *F. elaeidis* triggers more severe disease symptoms in *Aglaonema modestum* as compared with single infections (Zhang *et al*., 2022). This further highlight the difficulty to predict multispecies-complex dynamics, which are proposed to result from evolutionary competition and pathogen niche specialization (Abdullah *et al*., 2017).

Altogether these observations may indicate that the ability of taxa from the FHB complex to cooperate *vs*. compete may have a fitness cost which could be partially determined by the host. Such cost could for instance be associated with tradeoffs between the maintenance of virulence factors important to infect the host vs. antimicrobial factors important for competition with other taxa in the complex. Numerous studies showing that filamentous plant pathogens significantly impact the host microbiome, and recent discoveries suggesting that pathogen genomes encode large arrays of so-called effectors which are primarily targeted to outcompete other microbes (Flores-Nunez & Stukenbrock, 2024; Snelders et al., 2022). While at the same time, it was also shown that the maintenance of such large repertoire of virulence factors can also have an impact on pathogen fitness (Montarry et al., 2010). These observations are highly relevant for the cases where we saw possible signs of competition over plant immunity and plant resources in two cases where *F. graminearum* was combined with another *Fusarium* species or *Z. tritici*. This is also consistent with previous studies demonstrating that both *F. greaminearum* and *Z. tritici* have the capacity to manipulate the plant microbiome during infection, including the identification of specific effector proteins with antifungal activity in from *Z. tritici* (Kettles et al., 2018; Rojas et al., 2020). In fact, competition between *Fusarium* species as a strategy for biological control has been discussed and tested with some success (Aimé *et al*., 2013; Alabouvette *et al*., 2009).

We therefore conclude that the ability to reveal differences in the plant immune response to single vs. species infection is an important step forward in understanding the factors controlling FHB complex assembly, which further demonstrates the future potential of our experimental pipeline to be used a genetic tool for resistance screening.

## Data availability

All data is available within the manuscript or in the supplementary material. Additional information will be made available upon request to the corresponding author.

## Aknowledgment

We would like to acknowledge Fredric Hedlund and Ayano Tanaka for technical assistance with the growth facilities at SLU Biocentrum. We would also like to acknowledge the support of the Plant Protection Extensions ‘Växtskyddscentralerna’ of the Swedish Board of Agriculture ‘Jordbruksverket’ (https://jordbruksverket.se/) with field sampling across Sweden. We also thank Katarina Ihrmark and Maria Jonsson for their technical assistance.

